# High-fat diet induced loss of GABAergic inhibition decouples intrinsic and synaptic excitability in AgRP neurons

**DOI:** 10.1101/2021.05.31.446473

**Authors:** Austin C. Korgan, Wei Wei, Sophie L. A. Martin, Catherine C. Kaczorowski, Kristen M.S. O’Connell

**Author notes:** These authors contributed equally (Austin C. Korgan, Wei Wei).

## Abstract

Obesity is a progressive, relapsing disease with few therapies. Diet and lifestyle interventions are effective but are often temporary and many individuals regain weight. High-fat diet increases the excitability of AgRP neurons, a critical neuronal population for the regulation of food intake and body weight. Here we investigate the plasticity of AgRP neurons and the impact of high-fat diet on modulation by synaptic input. We find that diet-induced hyperexcitability of AgRP neurons is not reversed by a lower-fat diet intervention. High-fat diet is associated with changes in the synaptic modulation of AgRP neurons, with a paradoxical increase in inhibitory input accompanied by a loss of GABA-mediated inhibition due to a depolarizing shift in the reversal potential of the GABA-evoked Cl^−^ current. These findings reveal that high-fat diet leads to decoupling of intrinsic and synaptic excitability in AgRP neurons, such that hyperexcitability of AgRP neurons persists despite an increase in inhibitory input, revealing a mechanism for the difficulty in sustaining weight loss.

## Introduction

Despite substantial public health efforts, obesity prevalence in the United States has continued to rise, from ~30% in 2000 to more than 42% in 2018 (Hales, 2020). Beyond the social and cultural issues surrounding obesity and body weight, obesity carries with it serious medical implications and is associated with increased risk for and mortality from diseases such cardiovascular disease, type 2 diabetes, hypertension, osteoarthritis, cancer, Alzheimer’s disease, and Covid-19 (Abdelaal et al., 2017; Gao et al., 2020; Popkin et al., 2020; Profenno et al., 2010; Pugazhenthi et al., 2017). Obesity is the second leading cause of preventable death, accounting for nearly 20% of deaths in Americans aged 40-85 (Masters et al., 2013). Although the American Medical Association recognized obesity as a disease in 2013 (Kyle et al., 2016), there remain few safe, effective therapeutics for obesity (Van der Ploeg, 2000). Diet and lifestyle changes are an effective means of accomplishing weight loss, but the results are often temporary, with many individuals regaining lost weight within 5 years (Berk et al., 2018; Fothergill et al., 2016; Greenway, 2015; Hall and Guo, 2017; Sumithran and Proietto, 2013). This suggests that obesity, or obesity pathogenesis, is associated with a ‘resetting’ of the neural pathways that control energy homeostasis and body weight. Thus, successful, lasting interventions for obesity will need to address this remodeling, highlighting the importance of understanding how obesity and obesogenic factors such as a calorie-dense, high-fat diet impact the function of these pathways.

Located in the arcuate nucleus of the hypothalamus (ARH), AgRP neurons play a key role in the integration of peripheral and central signals to modulate food intake and body weight (Andrews et al., 2008; Garfield et al., 2016; Könner et al., 2007; Krashes et al., 2014; Morton and Schwartz, 2001; Varela and Horvath, 2012). Their activation is associated with hunger and food seeking (Baver et al., 2014; Mandelblat-Cerf et al., 2015; Takahashi and Cone, 2005; van den Top et al., 2004) and stimulation of AgRP neurons drives voracious feeding (Aponte et al., 2011; Chen et al., 2016; Krashes et al., 2011), while their inhibition or ablation results in hypophagia or starvation (Krashes et al., 2011; Luquet et al., 2005). Consumption of a high-fat diet (HFD) and obesity alter the function of AgRP neurons (Beutler et al., 2020; Mazzone et al., 2020), resulting in the persistent activation of these neurons, which become refractory to modulation by physiological cues of hunger or satiety (e.g., leptin) (Baver et al., 2014; Wei et al., 2015), thus, diet-induced dysfunction of AgRP neurons may represent a causal event in the pathogenesis of obesity.

There remain several unanswered questions with regard to the impact of body weight and diet on the function and modulation of AgRP neurons. First, AgRP neurons exhibit diet-induced plasticity in their intrinsic excitability following consumption of a high-fat diet (Baver et al., 2014; Wei et al., 2015), but it is unknown whether this plasticity is reversible and can be reset by a dietary intervention. Second, AgRP neurons receive abundant synaptic input and are rapidly modulated by the sensory detection of food (Betley et al., 2015; Chen et al., 2015; Mandelblat-Cerf et al., 2015), how the impact of diet and obesity impact synaptic modulation of AgRP neuronal excitability remains poorly understood. In this study, we set out to address both of these questions using *ex vivo* electrophysiology to directly assess the intrinsic and synaptic excitability of AgRP neurons from mice fed HFD for varying lengths of time. We find that diet-induced plasticity of AgRP neurons is not reversible and that the impact of HFD on AgRP neuronal excitability is long-lasting, even after acute HFD feeding. We also find that synaptic modulation of AgRP neurons is decoupled from intrinsic excitability; this is particularly evident in the action of GABAergic inputs, which no longer elicit an inhibitory effect in the postsynaptic AgRP neuron. Perforated patch clamp recordings demonstrate a loss of GABA-mediated inhibition in AgRP neurons, likely due to dyshomeostasis of intracellular Cl^−^. Our findings reveal a mechanistic explanation for the persistent effects of HFD and obesity and provide a mechanistic explanation for the difficulty many individuals experience with long-term weight loss.

## Methods

### Animals

The following transgenic strains were used in this study: *hrGFP-NPY* (JAX Stock #006417), C57Bl/6J (JAX Stock#000664), and *Vgat-IRES-Cre* (JAX Stock #028862). Founder mice were obtained from the JAX Repository and maintained by backcrossing with C57Bl/6J. The hrGFP-NPY mice were SNPtyped to confirm that the strain is on a congenic C57Bl/6J background except for the transgene insertion site on Chr7. All animal care and experimental procedures were approved by The Animal Care and Use Committee at The University of Tennessee Health Science Center and The Jackson Laboratory. Mice were maintained at 22-24° C on a 12h:12h light/dark cycle (0600 - 1800). All mice used for breeding were fed standard lab chow (UTHSC – Teklad 7912: 3.1 kcal/g metabolizable energy, 17 kcal% fat or JAX - LabDiets 5K0Q: 3.15 kcal/g metabolizable energy, 16.8 kcal% from fat). There was no significant effect of chow diet (Teklad v LabDiets) or site (UTHSC v JAX), so data from all control animals were combined. Mice were weaned at 21 days and group housed with same sex litter mates (n=2-5 mice/pen). At 8 weeks of age, offspring were randomly assigned to either standard chow or a high-fat diet (HFD; Research Diets D12451: 4.73 kcal/g metabolizable energy, 45 kcal% from fat) and maintained on the assigned diet for 8-10 weeks. The same HFD was used at both UTHSC and JAX and there was no significant effect of site on response to diet. Water was available *ad libitum*. Mice were weighed weekly and all mice were 10-18 weeks of age for all experiments.

### Electrophysiology

#### Slice preparation

For all experiments, brain slices were prepared between 0900 and 1030. Mice (10-18 weeks old) used for electrophysiology experiments were deeply anesthetized using isoflurane before decapitation and rapid removal of the brain. The brain was then submerged in ice-cold, oxygenated (95% O_2_/5% CO_2_) cutting solution (in mM: 119 NaCl, 90 sucrose, 2.5 KCl, 1 MgSO_4_, 2 CaCl_2_, 1.25 NaH_2_PO_4_, 23 NaHCO_3_, and 10 glucose). Coronal slices (250 μm) were cut using a vibratome (VT1000S, Leica) and incubated in oxygenated aCSF (in mM: 119 NaCl, 2.5 KCl, 1 MgSO_4_, 2 CaCl_2_, 1.25 NaH_2_PO_4_, 23 NaHCO_3_, and 10 glucose) for at least 1 h prior to recording.

#### Slice recording

Slices were transferred to a recording chamber constantly perfused (~2 ml/min) with oxygenated aCSF. GFP-positive AgRP/NPY neurons were identified using epifluorescence and standard GFP filters on a fixed-stage SliceScope 1000 microscope (Scientifica, Uckfield, UK) equipped with a digital camera (Q-Imaging, Surry, BC, Canada). All recordings were performed using a Multiclamp 700B amplifier and Digidata 1550A, controlled using Clampex 10.7 (Molecular Devices, San Jose, CA, USA). Data were digitized at 20 kHz and filtered at 5 kHz using the built-in four-pole Bessel filter of the Multiclamp 700B.

Recording pipettes were pulled from filamented thin-wall borosilicate glass (TW150F-4, World Precision Instruments) and had a resistance of 4-7 MΩ when filled with internal solution (for intrinsic excitability (AP) recordings, in mM: 130 K-gluc, 10 KCl, 0.3 CaCl_2_, 1 MgCl_2_, 1 EGTA, 3 MgATP, 0.3 NaGTP, and 10 HEPES, pH 7.35 with KOH; for synaptic recordings, in mM: 140 KCl, 0.3 CaCl_2_, 1 MgCl_2_, 1 EGTA, 3 MgATP, 0.3 NaGTP, and 10 HEPES, pH 7.35 with KOH; for perforated patch, Gramicidin A (Sigma) was dissolved in DMSO to a concentration of 5 mg/mL and then diluted in the synaptic internal solution (140 mM KCl) to a final concentration of 20 ug/mL). The liquid junction potential (LJP) between normal aCSF and the K-gluconate solution used for intrinsic recordings was +14.7 mV and was corrected. The LJP between aCSF and the KCl intracellular solution was +4.75mV and was not corrected.

Whole-cell current clamp recordings of resting membrane potential and spontaneous firing were recorded in the presence of DNQX (10 μM; Tocris) and picrotoxin (100 μM; Tocris). For experiments testing inhibition of AgRP neurons by leptin, mice were fasted overnight to promote intrinsic excitability and 100 nM leptin (National Hormone and Peptide Program) was bath applied. Whole-cell voltage-clamp recordings for mEPSC and mIPSC were conducted in the presence of TTX (1 μM; Tocris) and Picrotoxin (100 uM; Tocris) for mEPSCs and DNQX (10 μM; Tocris) for mIPSCs. We measured synaptic frequency, inter-event interval, amplitude, and τ-decay for mEPSC and mIPSC recordings.

Perforated patch voltage-clamp recordings for E_GABA_ were conducted in the presence of DNQX. Gigaohm seals were quickly established with target cells, then E_GABA_ measurements were made after the pipette resistance dropped to values between 40-80 MΩ, typically 15-45 min. Once resistance was stable, cells were voltage clamped at −70 mV, and +15 mV voltage steps were applied, ranging from −80 mV to 10 mV during which GABA (100 μM) was focally applied via pressure ejection using a Picospritzer III (Parker Hannifin, Hollis, NH USA) or PDES Pneumatic drug Ejection (npi electronic GmbH, Tamm, Germany). The I/V curve for each cell was calculated based on the current (pA) immediately prior to and following the GABA-puff.

Cell-attached recordings were conducted in voltage-clamp mode in the presence of picrotoxin (100 μM) for glutamate experiments and DNQX (10 μM) for GABA experiments. Normal aCSF was used as the intracellular solution. For both of these experiments, 10s puff of either glutamate (100 μM) or GABA (100 μM) were applied following the establishment of a loose-patch (15-75 MΩ).

### Data analysis and statistics

Post-synaptic current frequencies, amplitudes, inter-event intervals and decay time constants were measured using Clampfit 10.7 (Molecular Devices) and Axograph (AxoGraph, Inc). I/V curves from perforated patch recordings were calculated in Clampfit 10.7 (Molecular Devices). Statistical outliers were identified using the ROUT method (Q=1% cutoff threshold) as implemented in GraphPad Prism 9. Group differences were analyzed with two-way ANOVA followed by Tukey’s multiple comparisons *post hoc* test using Prism 8 and 9 (GraphPad). When required, three-way ANOVA was performed using the *aov* function in R (v3.6.3) using the R package r/emmeans for *post hoc* Tukey’s multiple comparison testing with Bonferroni’s correction. For repeated measures analysis, group differences were analyzed by two-way repeated measures-ANOVA using SPSS (IBM) or R. Data visualization was performed using Prism or r/ggplot2. For all statistical tests, a value of *p* < 0.05 was considered significant. Data are presented as the mean ± SEM; violin plots are presented as median ± quartile.

### Data Availability

All relevant data and analysis tools are available upon reasonable request from the authors.

## Results

### Female mice are resistant to diet-induced changes in AgRP neuronal hyperexcitability

To investigate the impact of sex and HFD on body weight, food consumption, AgRP neuronal excitability, and leptin sensitivity, we randomly assigned male and female 8-week-old C57Bl/6J mice to either a low-fat normal chow diet (NCD) or a high-fat/high-sugar diet (HFD). At 8 weeks of age, mice were randomly assigned to either NCD or HFD and maintained on the assigned diet for 8 weeks. Consistent with prior reports, male mice on HFD exhibit significant diet-induced weight gain, starting at ~2 weeks after HFD feeding and continuing throughout the 8-week experimental period (Fig 1A, *top*). In contrast, age-matched female mice fed HFD exhibit little weight gain over the 8-week HFD-feeding period, with no significant differences between the NCD- and HFD-fed groups at any time point, consistent with previous reports (Fig. 1A, *lower*) (Atamni et al., 2016; Dorfman et al., 2017; Hong et al., 2009; Hwang et al., 2010; Perez-Sieira et al., 2013; Pettersson et al., 2012; Salinero et al., 2018; Yakar et al., 2006).

**Figure 1:**
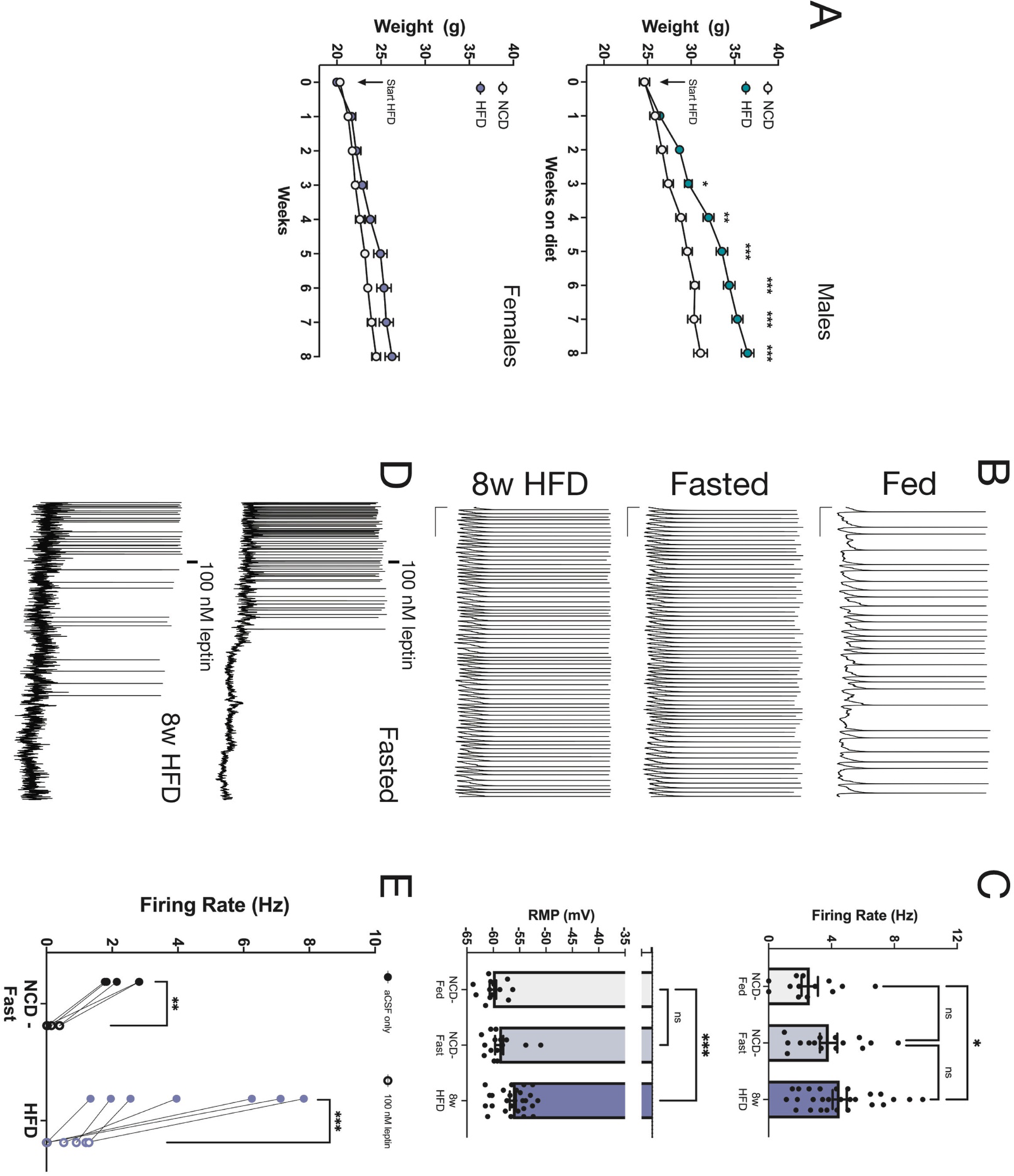
Female mice are resistant to diet-induced changes in AgRP neuronal hyperexcitability. **A.** Cumulative body weight curves for male (top) and female (bottom) mice during 8 weeks of HFD feeding. **B.** Representative traces of AgRP baseline firing rates from female mice fed NCD (top), following an overnight (16h) fast (middle), or 8 week HFD fed (bottom). **C.** 8 weeks of HFD feeding, but not fasting, significantly increase AgRP neuronal resting membrane potential (RMP) and firing rate. **D.** Representative traces of AgRP neuronal inhibition from female mice fed NCD (top) or HFD for 8 weeks (bottom). **E.** Leptin significantly inhibited AgRP neuronal firing in both NCD and 8 week HFD fed female mice. For all analysis, diet conditions compared to NCD using standard parametric ANOVA with a *post hoc* Tukey’s multiple comparisons test and a mixed-effects repeated measures ANOVA for repeated measures (* = p < 0.05; ** = p < 0.01; *** = p < 0.001).

We previously demonstrated that in male HFD-fed mice, AgRP neurons mice become hyperexcitable and refractive to modulation by physiological cues of hunger and satiety (Baver et al., 2014), with changes in neuronal function evident even after very brief periods of high-fat feeding, before the onset of significant weight gain (Wei et al., 2015). To determine if AgRP neurons from female mice undergo similar diet-induced remodeling, we used whole-cell patch clamp to measure the intrinsic properties of these neurons from female mice fed either NCD or HFD for 8 weeks (Fig 1B). In female mice, both overnight fasting (16h) and 8 weeks of HFD were associated with a slight increase in AgRP neuronal firing rate (Fig 1C, *top*; NCD-Fast: 3.82 ± 0.55, n = 15; HFD: 4.52 ± 0.45, n = 27, *F_(2, 52)_* = 3.33, p = 0.044), however while the absolute increase in neuronal firing due to fasting or HFD was comparable to what we previously observed in male mice (see (Baver et al., 2014; Wei et al., 2015), the relative increase in *f_AP_* was smaller in females due to a significant increase in the baseline firing rate in NCD-fed female mice compared to age-matched NCD-fed lean male mice (Females: 2.6 ± 0.52 s^−1^, n = 13; Males: 0.97 ± 0.31, n = 20; two-tailed *t*_38_ = 2.893, p = 0.007). As an additional measure of intrinsic excitability, we also measured the resting membrane potential (RMP), which we had previously reported to be depolarized by both fasting and HFD in male animals (Baver et al., 2014; Wei et al., 2015) and found that HFD, but not fasting, was associated with a significant depolarization of the RMP (Figure 1C, *lower*, NCD-fed = −60.0 ± 0.6 mV, n = 13; NCD-fast = − 58.9 ± 0.8 mV, n = 15; HFD: −56.3 ± 0.6 mV, n = 27, *f_(2,52)_* = 8.991, p = 0.0004).

Lastly, we assessed the leptin sensitivity of AgRP neurons from female mice following 8 weeks of HFD given that male mice fed a HFD for 8 weeks develop hypothalamic leptin resistance (Baver et al., 2014; El-Haschimi et al., 2000; Münzberg et al., 2004). We determined the impact of HFD on the leptin-induced inhibition of AgRP neurons from female mice as in (Baver et al., 2014): mice were fasted for ~16h (to maximize AgRP neuronal activity) and slices prepared for whole-cell current clamp recording of AgRP firing. As shown in Figure 1D and E, bath application of 100 nM leptin significantly and robustly inhibited AgRP neuronal activity in brain slices from fasted NCD female mice (NCD: 2.3 ± 0.2 s^−1^; +leptin: 0.2 ± 0.1 s^−1^, n = 5, paired *t_5_* = 9.204, *p* = 0.002). In contrast to our previous findings in male mice, AgRP neurons from female mice on HFD remained highly sensitive to leptin (HFD: 4.4 ± 0.6 s^−1^; +leptin: 0.6 ± 0.2 s^−1^, n = 6, paired *t_6_* = 3.73, p = 0.01). Taken together, these data suggest that there is strong sexual dimorphism in both the baseline function of AgRP neurons in female mice and in the neuronal response to HFD, with female mice exhibiting resistance to diet-induced weight gain and modestly increased neuronal activity independent of body weight.

### HFD-induced hyperexcitability of AgRP neurons is not reversed by refeeding with a low-fat NCD

We previously demonstrated that consumption of HFD, even for brief periods, is associated with hyperexcitability of AgRP neurons and resistance to modulation by physiological cues such as hunger and leptin (Baver et al., 2014; Wei et al., 2015). To determine the persistence of this change in excitability and assess whether ‘normal’ excitability can be restored, which is likely key for the development of effective weight-loss therapies, we modeled a common, translationally relevant dietary intervention – switching to a lower-fat, lower-calorie diet. Since we did not observe significant weight gain, leptin resistance, or remodeling of AgRP neuronal excitability in HFD-fed female mice, only male mice were used for these experiments. Male mice were fed a HFD for 8 weeks to induce DIO, then switched to a low-fat NCD for 10 weeks (Fig. 2A); age-matched control animals were maintained on NCD for the entire 18-week period. As shown in Fig. 2B and C, 10 weeks of lower calorie NCD feeding following 8 weeks of HFD failed to restore the excitability of AgRP neurons to the level observed in NCD-only mice (NCD_only_: 1.1 ± 0.3 s^−1^, n = 21; HFD→NCD: 3.8 ± 0.5 s^−1^, n = 25; *t_44_* = 3.929, p = 0.0003). In fact, the intrinsic firing rate of AgRP neurons in the HFD→NCD mice was nearly identical to that measured in HFD only mice (HFD_only_: 3.4 ± 0.4 s^−1^, n = 27; *t_50_* = 0.4564, p = 0.65, secondary analysis of HFD data from (Baver et al., 2014)).

**Figure 2:**
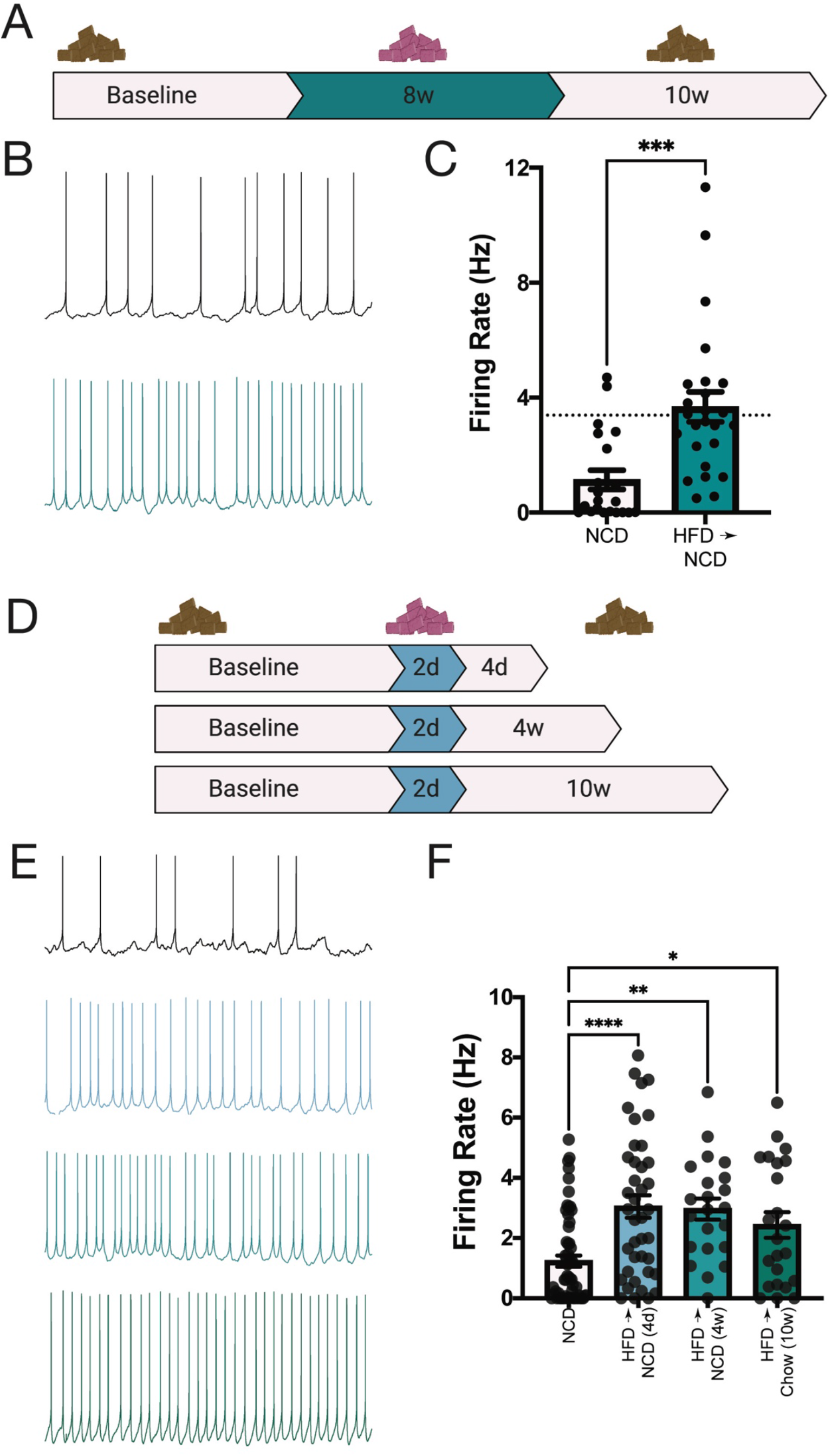
HFD-induced hyperexcitability of AgRP neurons is not reversed by refeeding with a low-fat NCD. **A.** Timeline for NCD refeeding experiments following 8 weeks of HFD feeding. **B.** Representative traces of AgRP baseline firing rates from male mice maintained on NCD (top) or treated with NCD refeeding following 8 week HFD feeding (bottom). **C.** AgRP neuronal firing rate remained significantly elevated following 10 weeks of NCD feeding and was similar to previously reported baseline AgRP neuronal firing rates HFD fed male mice (dashed line). **D.** Timeline for NCD refeeding experiments following 4 days of HFD feeding. **E.** Representative traces of AgRP baseline firing rates from male mice maintained on NCD (top) or fed HFD for 2 days and treated with NCD refeeding for 4 days (top middle), 4 weeks (bottom middle), or 10 weeks (bottom). **F.** AgRP neuronal firing rates remained significantly elevated by HFD_2d_ followed by 4 day, 4 week, and 10 week NCD refeed compared to NCD fed mice. For all analysis, diet conditions compared to NCD using standard parametric ANOVA with a *post hoc* Tukey’s multiple comparisons test (* = p < 0.05; ** = p < 0.01; *** = p < 0.001, **** = p<0.0001).

Previously, we observed that even very brief periods of HFD feeding (<1 week) were sufficient to promote hyperexcitability of AgRP neurons, though these neurons are still sensitive to inhibition by leptin sensitivity (Wei et al., 2015). Thus we hypothesized that AgRP neurons might retain plasticity and more amenable to a dietary intervention at this time point. To test this, we fed male mice HFD for 4 days, then switched them to NCD for either 4 days, 4 weeks, or 10 weeks (Fig. 2D) before recording AgRP neuron intrinsic excitability in brain slices containing ARH. As shown in Fig. 2E and F, although none of the NCD feeding paradigms were able to fully reverse the effect of HFD on neuronal excitability, increasing duration of NCD feeding post-HFD is associated with a decrease in the intrinsic firing rate of AgRP neurons (NCD_only_: 1.2 ± 0.19 s^−1^, n = 60; HFD→NCD_4d_: 3.1 ± 0.38^−1^, n = 40; HFD→NCD_4w_: 3.0 ± 0.35^−1^, n = 22; HFD→NCD_10w_: 2.4 ± 0.43^−1^; *F_(3,141)_ =* 9.458, p < 0.0001).

### Brief periods of HFD feeding influence synaptic excitability in AgRP neurons in a sexually dimorphic manner

Overnight fasting increases the intrinsic excitability of AgRP neurons (Baver et al., 2014; Liu et al., 2012; Takahashi and Cone, 2005; Yang et al., 2011), an effect that requires a corresponding change in excitatory glutamatergic input (Liu et al., 2012; Yang et al., 2011), raising the possibility that the increase in neuronal excitability following 2 days of HFD feeding (Wei et al., 2015) is associated with an increase in excitatory synaptic input. We therefore determined the impact of 2d of HFD (HFD_2d_) on excitatory input using whole-cell patch clamp to record the frequency and amplitude of miniature excitatory postsynaptic currents (mEPSC) in AgRP neurons from age-matched male and female NPY-GFP mice randomly assigned to either HFD for 2d or maintained on NCD. The impact of HFD on synaptic excitability in AgRP neurons is poorly understood in either sex, so both male and female mice were included in these experiments.

Consistent with our observation that AgRP neurons from female NCD mice exhibit a significantly higher baseline firing rate than age- and diet-matched male mice, there was a significant main effect of sex on the frequency of mEPSCs in AgRP neurons as baseline (*f*_mEPSC_). Neurons from NCD female mice exhibited a significantly higher *f*_mEPSC_ compared to those measured in AgRP neurons from NCD male mice (Figure 3, A and B; NCD Females: 4.5 ± 0.5, n = 10; NCD Males: 2.6 ± 0.3, n = 29, *F_(1,94)_* (Sex) = 11.60, p = 0.0001). There was also a significant main effect of diet on the f_mEPSC_ (*F_(1,94)_* = 15.22, p = 0.0002) as the frequency of excitatory input was increased in both HFD_2d_ males and females (HFD_2d_ Females: 5.6 ± 0.04, n = 22; HFD_2d_ Males: 4.7 ± 0.4, n = 28). However, although the absolute *f*_mEPSC_ is similar between the male and female HFD_2d_ groups, *post hoc* pairwise comparisons indicated that this increase was significant only in AgRP neurons from male mice (Males, adj. p = 0.0004; Females, adj. p = 0.14), again likely because the baseline *f*_mEPSC_ is higher in neurons from NCD female mice, suggesting that AgRP neurons from female mice are generally more excitable than those from males.

**Figure 3.**
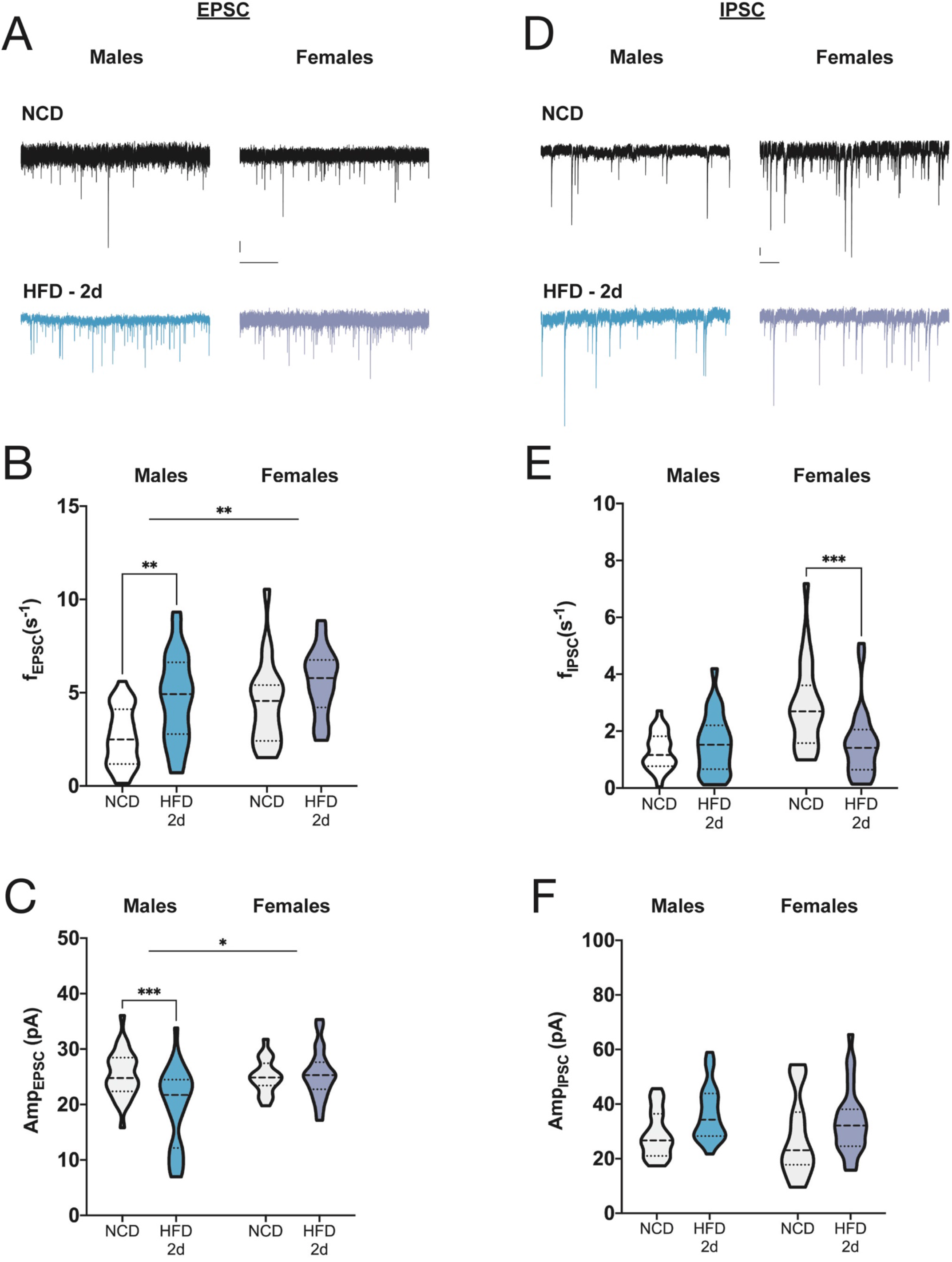
Brief periods of HFD feeding influence synaptic excitability in AgRP neurons in a sexually dimorphic manner. **A.** Representative traces of mEPSCs onto AgRP neurons from male mice fed NCD or HFD for 2d. **B.** Representative traces of mEPSCs onto AgRP neurons from female mice fed NDC or HFD for 2d. **C.** Mean mEPSC frequency in AgRP neurons from NCD and HFD_2d_ male and female mice (males: NCD n = 29; HFD_2d_ n = 29; females: NCD n = 19; HFD_2d_ = 22). **D.** Mean mEPSC amplitude in AgRP neurons. **E.** Mean mIPSC frequency in AgRP from NCD and HFD_2d_ male and female mice (males: NCD n = 31; HFD_2d_ = 25; females: NCD n = 24; HFD_2d_ n = 29). **F.** Mean mIPSC amplitude in AgRP neurons. For all violin plots, dashed line indicates median, dotted lines indicate quartiles. Statistical comparisons using two-way ANOVA with sex and diet as main effects; post hoc pairwise comparisons performed using Tukey’s multiple comparisons test (* = p < 0.05; ** = p < 0.01; *** = p < 0.001, **** = p<0.0001).

Although the amplitude of the mEPSC was not different in AgRP neurons from NCD male and female mice (Figure 3C; NCD Females: 25.08 ± 0.7 pA; NCD Males: 25.41 ± 0.78 pA, adj. p > 0.99), there was a significant interaction between sex and diet (*F_(1,94)_* = 8.097, p = 0.005), as the amplitude of the mEPSC in AgRP neurons from male HFD_2d_ mice was significantly decreased compared to that measured in NCD controls, while diet had no effect on mEPSC amplitude in neurons from female mice (HFD_2d_ Males: 19.81 ± 1.3 pA, adj. p = 0.0001 vs NCD; HFD_2d_ Females: 25.41 ± 0.96, adj. p = 0.97 vs NCD). A change in the frequency of synaptic events is usually interpreted as an alteration in presynaptic function (e.g., increased release probability or synapse number), while changes in amplitude are thought to reflect primarily changes in postsynaptic mechanisms, these results suggest that in male, but not female, mice, brief HFD feeding alters AgRP synaptic excitability via both pre- and postsynaptic mechanisms.

We also investigated the influence of 2d of HFD feeding on inhibitory GABAergic input to AgRP neurons by measuring the miniature inhibitory postsynaptic currents (mIPSC) in ARH-containing brain slices from age-matched male and female mice fed either NCD or HFD for 2d. There was a significant main effect of sex on the *f*_mIPSC_ (*F_(1,98)_* = 12.82, p = 0.0005), reflecting an increase in the *f*_mIPSC_ in AgRP neurons from NCD female mice compared to NCD male mice (Figure 3, A, D, and E; NCD Females: 2.89 ± 0.3, n = 24; NCD Males: 1.3 ± 0.1, n = 28). There was a significant interaction between sex and HFD_2d_ on *f*_mIPSC_ (Sex:Diet *F_(1,98)_* = 12.39, p = 0.0007), as HFD_2d_ significantly decreased *f*_mIPSC_ in neurons from female mice (HFD_2d_: 1.57 ± 0.2, adj. p = 0.0005 vs NCD females), but had no effect on the *f*_mIPSC_ in neurons from male HFD_2d_ mice (Figure 3E; HFD_2d_: 1.56 ± 0.2, adj. p > 0.99 vs NCD Males). There was no main effect of sex on the mIPSC amplitude (Figure 3F), but 2d of HFD feeding was associated with a significant decrease in the amplitude of the mIPSC (Diet: *F_(1,98)_* = 9.156, p = 0.003). This effect was driven by a significant increase in the mIPSC amplitude in AgRP neurons from both male (NCD: 28.8 ± 1.5 pA; HFD_2d_ = 36.6 ± 2.1, adj. p = 0.03), but not in those from female mice (NCD: 28.07 ± 3.0; HFD_2d_: 33.7 ± 2.2, adj. p = 0.15).

### Long-term HFD does not alter excitatory synaptic input to AgRP neurons

Given the persistent hyperexcitability of AgRP neurons following HFD feeding, we next determined whether the diet-induced synaptic plasticity observed after only 2d of HFD is maintained with longer periods on HFD, such as those associated with the development of diet-induced obesity (DIO). Age-matched male and female NPY-GFP mice were randomly assigned to either NCD or HFD at 8 weeks of age and maintained on the assigned diet for 8 weeks (Figure 1A). To determine the impact of diet on metabolic-state dependent synaptic plasticity (Liu et al., 2012; Wang et al., 2021; Yang et al., 2011), a subset of mice from both the NCD and HFD groups were fasted for 16h prior to whole-cell recording of the mEPSC in AgRP neurons. Consistent with previous reports (Liu et al., 2012; Yang et al., 2011), we observed a significant increase in the *f*_mEPSC_ in AgRP neurons from male NCD mice (Figure 4, A and C; NCD_fed_: 3.01 ± 0.6; NCD_fast_: 7.04 ± 0.6, adj. p = 0.0006). There was no main effect of sex on the *f*_mEPSC_ (Sex: *F_(1, 181)_* = 2.43, p = 0.12) as AgRP neurons from female mice exhibited a similarly robust response to fasting (NCD_fed_: 2.7 ± −.4; NCD_fast_: 6.8 ± 0.6, adj. p < 0.0001). Despite the increased intrinsic excitability of AgRP neurons under these conditions (Baver et al., 2014), 8w of HFD feeding did not significantly alter the baseline *f*_mEPSC_ in neurons from fed male and female mice (Figure 4, A-C; Females HFD_fed:_ 4.0 ± 0.5, n = 26; Males HFD_fed_: 2.3 ± 0.2, n = 18, Diet: *F_(1,181)_* = 1.011, p = 0.31), nor did it affect the response to fasting, as *f*_mEPSC_ was significantly increased in both males and females following an overnight fast to a similar extent as observed in neurons from NCD_fast_ animals (Figure 4, A-D; Females HFD_fast_: 7.2 ± 1.0, n = 25; Males HFD_fast_: 5.2 ± 0.7, n = 28, Metab.State: *F_(1,181)_* = 62. 627, p = 2.27 x 10^−13^).

**Figure 4.**
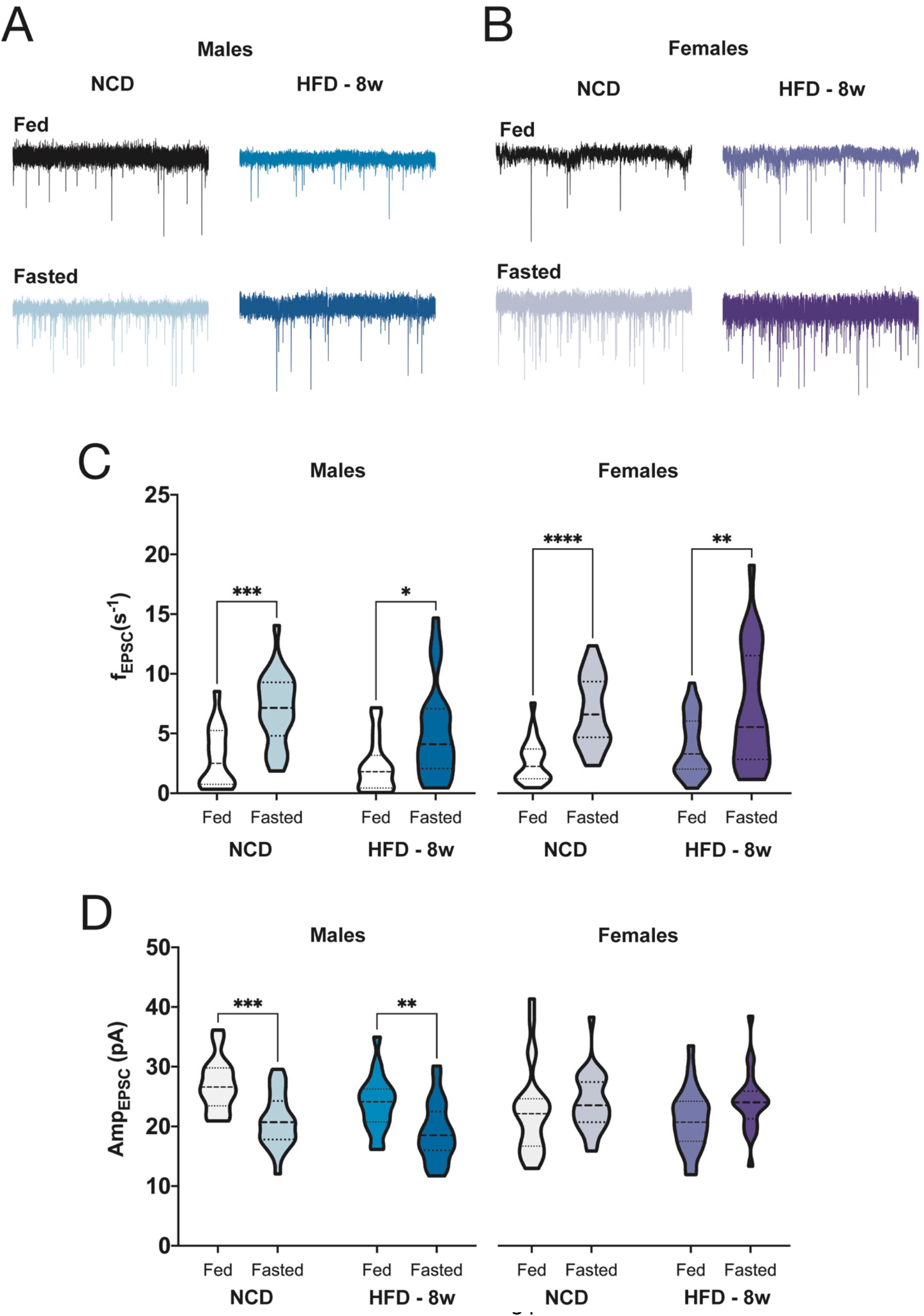
Long-term HFD does not alter excitatory synaptic input to AgRP neurons. **A.** Representative traces of mEPSCs onto AgRP neurons in fed or fasted male mice on either NCD or HFD_8w_. **B.** Representative traces of mEPSCs onto AgRP neurons in fed or fasted female mice on either NCD or HFD_8w_. **C.** Mean mEPSC frequency onto AgRP neurons from fed and fasted male and female mice fed either NCD or HFD_8w_ (males: n = 16 – 28 neurons/group; females: n = 24 – 27 neurons/group). **D.** Mean mEPSC amplitude in AgRP neurons. For all violin plots, dashed line indicates median, dotted lines indicate quartiles. Statistical comparisons using three-way ANOVA with sex, diet, and metabolic state as main effects; post hoc pairwise comparisons performed using Tukey’s multiple comparisons test (* = p < 0.05; ** = p < 0.01; *** = p < 0.001, **** = p<0.0001).

When we investigated the impact of long-term HFD feeding on the amplitude of the mEPSC, we found a significant interaction between sex and metabolic state (Sex:State: *F_(1,181)_* = 25.493, p = 1.1 x 10^−6^). In AgRP neurons from male mice, fasting was associated with a significant decrease in the amplitude of the mEPSC, regardless of diet (Figure 4D; NCD_fed_: 27.0 ± 1.1 pA; NCD_fast_: 21.0 ± 0.9 pA, adj. p = 0.0003; HFD_fed_: 23.9 ± 1.1 pA; HFD_fast_ = 18.95 ± 0.9 pA, adj. p = 0.0015). However, in AgRP neurons from female NCD mice, fasting had no effect on the mEPSC amplitude (Figure 4D; NCD_fed_: 22.6 ± 1.5 pA; NCD_fast_: 24.05 ± 0.9 pA, adj. p = 0.32) and although HFD_8w_ did not alter the amplitude of the mEPSC under baseline HFD_fed_ conditions, it was associated with a small, but significant increase in the mEPSC amplitude (Figure 4D; HFD_fed_: 21.1 ± 1.0 pA; HFD_fast_: 24.2 ± 1.0, adj. p = 0.03).Taken together, these results suggest that neither HFD nor obesity impact excitatory glutamatergic presynaptic input to AgRP neurons or metabolic state-dependent synaptic plasticity, but may influence the postsynaptic response to excitatory input.

### Long-term HFD feeding alters inhibitory synaptic input to AgRP neurons

Since there was no significant effect of 8 week HFD diet on excitatory glutamatergic input to AgRP neurons (Figure 4), we reasoned that diet-induced hyperexcitability of AgRP neurons might be due, at least in part, to a disruption of excitation/inhibition (E/I) balance. AgRP neurons are rapidly inhibited by the sensory detection of food (Betley et al., 2015; Chen et al., 2015; Mandelblat-Cerf et al., 2015), a response that is blunted in mice fed a HFD (Beutler et al., 2020; Mazzone et al., 2020), suggesting that 1) inhibitory input is a key factor in the physiological regulation of AgRP neurons and 2) HFD and/or DIO alters this input. We therefore hypothesized that long-term consumption of a HFD might be associated with an overall loss of GABAergic input to AgRP neurons. To test this, we recorded miniature inhibitory postsynaptic currents (mIPSC) in AgRP neurons from age-matched male and female NPY-GFP mice randomly assigned either NCD or HFD for 8 weeks.

As shown in Figure 5, in AgRP neurons from male mice, *f*_mIPSC_ is remarkably low, regardless of metabolic state (NCD_fed_: 0.53 ± 0.07 s^−1^, n = 29; NCD_fast_: 0.72 ± 0.1, n = 29, adj. p = 0.7). However, there is a significant interaction between sex and metabolic state (Sex:State *F_(1,231)_* = 8.662, p = 0.004) and *f*_mIPSC_ is significantly increased in AgRP neurons from NCD_fed_ female mice (NCD_fed_: 3.4 ± 0.3 s^−1^, n = 35) relative to both male NCD_fed_ (adj. p = 9.4 x 10^−9^ vs NCD_fed_ males) and female NCD_fast_ (NCD_fast_: 1.4 ± 0.2 s^−1^, n = 35, adj. p = 1.5 x 10^−5^ vs NCD_fed_ females). In neurons from both male and female mice, there was a significant main effect of diet on *f*_mIPSC_ in AgRP neurons (Diet: *F_(1, 231)_* = 43.613, p = 2.6 x 10^−10^), but the direction of the effect was opposite of what we predicted based on the patterns of spontaneous firing in AgRP neurons from both male and female mice after HFD (Figure 1, C-E and (Baver et al., 2014; Wei et al., 2015)). Indeed, long-term consumption of HFD was associated with an increase in the *f*_mIPSC_ in AgRP neurons from male mice (Figure 5, A and C; Males HFD_fed_: 3.0 ± 0.5 s^−1^, n = 29, adj. p = 1.37 x 10^−6^ vs NCD_fed_) that was not altered by metabolic state (Figure 5C; HFD_fast_: 2.4 ± 0.3 s^−1^, adj. p = 0.2 vs HFD_fed_). In neurons from female HFD_fed_ mice, there was a trend of the *f*_mIPSC_ to increase over NCD controls but this was not significant (Figure 5, B and C; HFD_fed_: 4.2 ± 0.8 s^−1^, n = 20, adj. p = 0.1). The effect of metabolic state on inhibitory input to AgRP neurons was blunted in neurons from female mice on HFD, as *f*_mIPSC_ in HFD_fast_ neurons was significantly greater than NCD_fast_ controls (Figure 5D; HFD_fast_: 3.1 ± 0.4 s^−1^, n = 28, adj. p = 0.0004) and was not significantly different form HFD_fed_ (adj. p = 0.051).

**Figure 5:**
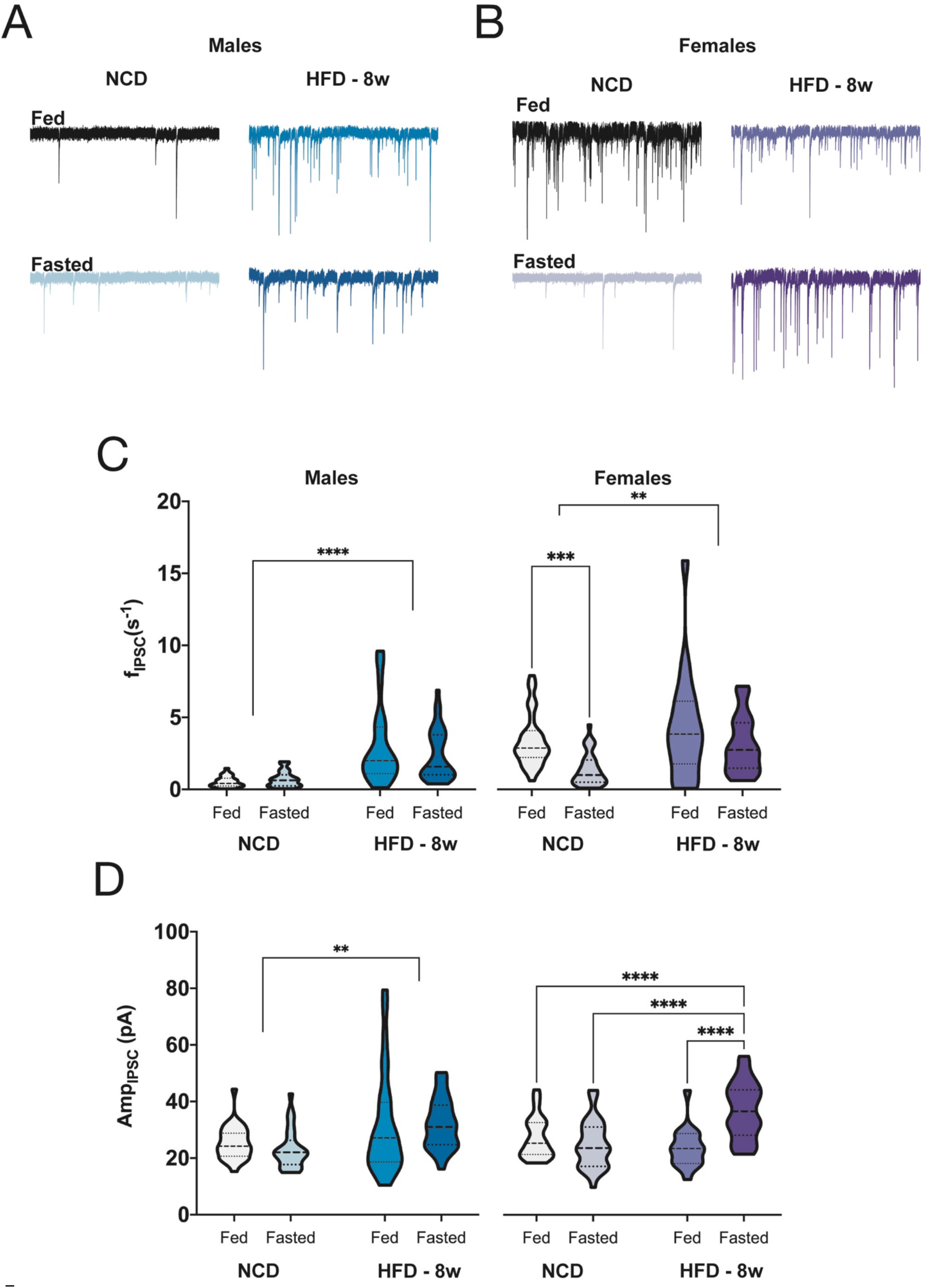
Long-term HFD feeding alters inhibitory synaptic input to AgRP neurons. **A.** Representative traces of mIPSCs onto AgRP neurons in male mice fed NCD or HFD_8w_. **B.** Representative traces of mIPSCs onto AgRP neurons in female mice fed NCD or HFD_8w_. **C.** Mean mIPSC frequency onto AgRP neurons (males: n = 29 – 35 neurons/group). **D.** Mean mIPSC amplitude from AgRP neurons. For all violin plots, dashed line indicates median, dotten lines indicate quartiles. Statistical comparisons used three-way ANOVA with sex, diet, and metabolic state and main effects with a *post hoc* Tukey’s multiple comparisons test (* = p < 0.05; ** = p < 0.01; *** = p < 0.001, **** = p<0.0001).

We also observed a significant effect of sex, diet, and metabolic state on the amplitude of the mIPSC (Sex:Diet:State *F_(3, 221)_* = 3.844, p = 0.01), such that in AgRP neurons from male mice, the mIPSC amplitude was increased in neurons from HFD_8w_ mice, regardless of metabolic state (NCD_fed_: 25.4 ± 1.1 pA; HFD_fed_: 31.61 ± 3.3 pA, adj. p = 0.01; NCD_fast_: 23.1 ± 1.3 pA; HFD_fast_: 32.19 ± 1.5 pA, adj. p = 0.0003). By comparison, mIPSC amplitude in neurons from female mice was comparable in both NCD_fed_ (27.2 ± 1.2 pA) and HFD_fed_ (23.7 ± 1.7 pA, adj. p = 0.22), but was significantly increased after fasting only in HFD mice (NCD_fast_: 25.01 ± 1.4 pA; HFD_fast_ = 36.9 ± 1.9, adj. p = 3.1 x 10^−6^).

### Long-term HFD feeding is associated with depolarization of E_GABA_, resulting in a loss of GABA-mediated synaptic inhibition

Our data suggest that although AgRP neurons receive both excitatory and inhibitory synaptic input, the postsynaptic response to these inputs is decoupled from intrinsic excitability following long-term HFD consumption in both male and female mice. To further confirm this finding, we specifically evaluated the impact of glutamatergic and GABAergic signaling on the spontaneous firing of AgRP neurons from NCD and HFD_8w_ mice.

Our results suggest that the modulation of glutamatergic presynaptic input is unaffected by HFD feeding but that the postsynaptic response to this input may be altered (Figure 4), raising the possibility that AgRP neurons may fail to respond appropriately to glutamatergic signaling. To test whether glutamate can still stimulate firing of AgRP neurons, we recorded the spontaneous firing rate of AgRP neurons from NCD_fed_ and HFD_fed_ mice in response to a brief (1 s) puff of 10 μM glutamate. These experiments were performed using the cell-attached configuration of the patch clamp technique to maintain intracellular ion concentrations at their physiological gradients. As shown in Figure 6, A and B, there was no significant effect of diet on the response to direct glutamate stimulation (Diet: *F_(1,22)_* = 0.48, p = 0.5), as neurons from both NCD and HFD mice responded with a significant increase in the firing rate (NCD_fed_: Baseline = 0.71 ± 0.22 s^−1^, +Glut = 5.24 ± 1.2 s^−1^; HFD_fed_: Baseline = 1.4 ± 0.5 s^−1^, +Glut = 6.4 ± 2.3 s^−1^; *F_(1,22)_* = 14.32, p = 0.0001).

**Figure 6:**
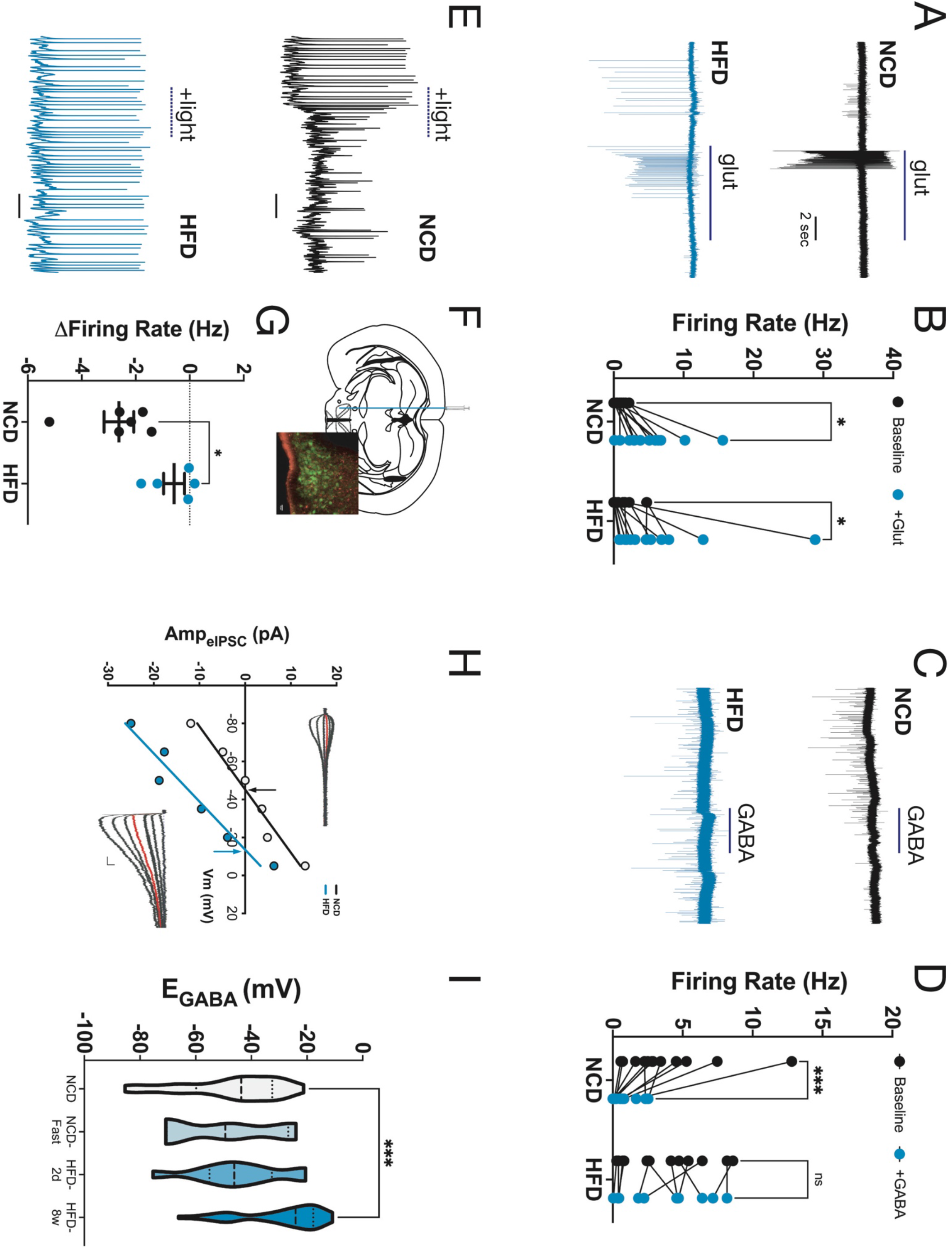
Long-term HFD feeding is associated with depolarization of E_GABA_, resulting in a loss of GABA-mediated synaptic inhibition. **A,** Representative traces of the response to a 1s puff of 10 μM glutamate in AgRP neurons from NCD (top) and HFD (bottom) AgRP neurons. All recording made using the loose-patch cell-attached configuration. **B.** AgRP neuronal firing rates before (black dots) and after (blue dots) puff application of glutamate (NCD: n = 12 neurons, HFD: n = 12 neurons; repeated measures two-way ANOVA with Tukey’s *post hoc* multiple comparisons test). **C.** Representative traces illustrating the response of AgRP neurons from NCD (top) and HFD (bottom) mice to puff application of 100 μM GABA. **D.** AgRP neuronal firing rates before (black dots) and after (blue dots) GABA application (NCD: n = 12 neurons; HF = 12 neurons, repeated measures two-way ANOVA with Tukey’s *post hoc* multiple comparisons test). **E.** Representative traces from optogenetically inhibited AgRP neurons from NCD (top) and HFD (bottom) mice. **F.** ChR2 was expressed in presynaptic inhibitory Lepr+ neurons from the ventral dorsomedial hypothalamus (vDMH) (top) (inset, terminals in red). **G.** Change in AgRP neuronal firing rate in response to light-evoked activation of GABAergic vDMH presynaptic terminals (Light – Baseline; paired t-test). **H.** Representative I/V curves show E_GABA_ is depolarized by approximately +20 mV in HFD males compared to NCD (inset; representative traces from NCD (top left) and HFD (bottom right) males. Red traces = −40 mV). **I.** Mean E_GABA_ in AgRP neurons from NCD (n = 24 neurons), NCD_fast_ (n = 7 neurons), HFD_2d_ (n = 10 neurons), and HFD_8w_ (n = 37 neurons) mice. Statistical comparisons using ordinary one-way ANOVA with Dunnett’s *post hoc* multiple comparisons test. (* = p < 0.05; ** = p < 0.01; *** = p < 0.001, **** = p<0.0001).

In contrast, when we performed a similar experiment using GABA to test the inhibitory response, we found a significant effect of diet on the inhibition of spontaneous firing (Diet: *F_(1,22)_* = 7.589, p = 0.02). In neurons from NCD_fast_ mice, puff application of 100 μM GABA significantly decreased the firing rate (Figure 6, C and D; NCD_fast_: Baseline = 3.9 ± 0.98, +GABA = 0.76 ± 0.3, adj. p = 0.0005), while in neurons from HFD_fed_ mice, GABA had no effect on the firing rate of AgRP neurons (Figure 6, C and D; Baseline = 3.7 ± 0.9, +GABA = 3.4 ± 0.8, adj. p = 0.9). In fact, in 25% of HFD neurons (3 of 12), GABA application resulted in an increase in the firing rate (Figure 6D), strongly suggesting that the intracellular Cl^−^ gradient is disrupted by HFD in AgRP neurons.

To confirm that AgRP neurons from HFD mice fail to be inhibited by GABA in a physiological context, we next used optogenetics to determine whether activation of a defined GABAergic projection to AgRP neurons was still able to modulate firing of the postsynaptic AgRP neuron. To this end, we expressed ChR2 in GABAergic Lepr^+^ neurons in the ventral dorsomedial hypothalamus (vDMH), a population that has been demonstrated to directly inhibit the firing of AgRP neurons and suppress food intake (Garfield et al., 2016) (Figure 6F). We used whole-cell patch clamp to monitor the spontaneous firing of AgRP neurons during photostimulation of ChR2-mCherry^+^ terminals from vDMH^Lepr^ neurons in the ARH. As shown in Figure 6E and G, in brain slices from NCD^fast^ mice, photoactivation of vDMH terminals rapidly inhibited firing in the postsynaptic AgRP neurons (Fig. 6G, Δ firing = −2.62 ± 0.55), but in the ARH of HFD_fed_ mice, photostimulation of ChR2+ vDMH terminals had no significant effect on the firing rate of AgRP neurons (Fig. 6G, Δ firing = −0.58 ± 0.4).

In adult neurons, GABA functions as an inhibitory neurotransmitter because intracellular Cl^−^ concentrations are maintained at a very low level due to the activity of the K^+^-Cl^−^ cotransporter KCC2 (*Slc12a5*), resulting in a hyperpolarized reversal potential for the GABA-evoked Cl^−^ current (E_GABA_) through GABA_A_-R, which are ligand-gated Cl^−^ channels. We therefore hypothesized that HFD leads to dysfunction in the activity of KCC2, leading to dissipation of the neuronal Cl^−^ gradient and a loss of GABAergic inhibition. However, KCC2 is an electroneutral transporter, precluding direct electrophysiological assessment of its function. Thus, we indirectly assessed KCC2 function and the Cl^−^ gradient in AgRP neurons by measuring E_GABA_. Because conventional whole-cell patch clamp necessarily disrupts the intracellular Cl^−^ concentration, we used the gramicidin perforated patch method for these experiments. As shown in Figure 6, H and I, E_GABA_ in AgRP neurons from NCD_fed_ mice was −47.09 ± 3.6 mV. There was no significant shift in E_GABA_ in AgRP neurons from either NCD_fast_ (E_GABA_ = −51.01 ± 3.6 mV, adj. p = 0.9 v NCD_fed_) or HFD_2d_ (E_GABA_ = − 44.4 ± 5.2 mV, adj. p = 0.96 vs NCD_fed_). However, consistent with our observation that GABA no longer inhibits AgRP neurons from HFD_8w_ mice, there was a significant depolarizing shift in the GABA reversal potential (E_GABA_ = −29.4 ± 2.4 mV, adj. p = 0.0003 vs NCD_fed_).

## Discussion

Obesity has been a significant public health concern for the past 50 years and is a significant comorbidity for diseases such as type 2 diabetes, cardiovascular disease, cancer, neurodegenerative diseases such as Alzheimer’s disease, and COVID-19. Despite decades of research, safe, effective, and lasting therapeutics are still lacking. Diet and lifestyle interventions are a generally effective approach for weight loss, but long-term adherence is generally low and most individuals regain the lost weight within 5 years (Foster et al., 2010; Hall and Guo, 2017; Nordmo et al., 2020; Webb and Wadden, 2017). Evidence from our lab (Baver et al., 2014; Wei et al., 2015) and others (Beutler et al., 2020; Mazzone et al., 2020; Rossi et al., 2019) indicates that high-fat, calorie-dense diets cause remodeling of the neurons and circuits that govern energy balance, highlighting a critical role for CNS plasticity in the development and maintenance of the obese state. Despite this, our understanding of how diet and obesity alter CNS function remains relatively poor. In this study, we aimed to fill that gap by investigating the impact of a high-fat diet on the intrinsic and synaptic excitability of AgRP neurons from both male and female mice. Our results reveal that AgRP neuronal function is sexually dimorphic, that the diet-induced hyperexcitability of AgRP neurons persists even after switching to a lower fat diet, and that diet is associated with a decoupling of intrinsic and synaptic excitability in AgRP neurons, most notably resulting in depolarization of E_GABA_ and decreased efficacy of GABA as an inhibitory neurotransmitter.

### AgRP neuronal function and response to HFD is sexually dimorphic

The hypothalamus is a sexually dimorphic area (Spring et al., 2007) and it has been previously reported that the response to obesogenic diets differs between males and females (Dorfman et al., 2017; Palmer and Clegg, 2015; Perez-Sieira et al., 2013; Pettersson et al., 2012; Salinero et al., 2018). However, the inclusion of females in mechanistic studies of hypothalamic function remains low and to our knowledge, there is very little, if anything, known about how diet or metabolic state influences the function of AgRP neurons in females. AgRP neurons are involved in the modulation of a variety of female-specific behaviors, such as fertility (Egan et al., 2017; Padilla et al., 2017), nest-building (Li et al., 2019), and lactation (Phillips and Palmiter, 2008), emphasizing the importance of understanding how diet and obesity influences the intrinsic and synaptic excitability of AgRP neurons in female mice.

Using whole-cell patch clamp recording, we found that at baseline, AgRP neurons from lean, NCD female mice are generally more excitable than neurons from male mice (Figure 1B and C). AgRP neurons from female mice also retained their sensitivity to inhibition by leptin as evidenced by the robust inhibition of AgRP neurons by leptin even after 8 weeks of HFD feeding, a paradigm that was associated with leptin resistance in male mice (Baver et al., 2014). Interestingly, the absolute magnitude of the increase in intrinsic excitability in female mice was comparable to that measured in male mice (Figure 1C, (Baver et al., 2014)), but since neurons from female animal have a higher baseline, the relative change is much smaller, which may contribute to the resistance to diet-induced weight gain in female mice over this time frame. Metabolism and substrate preference are sexually dimorphic, with females tending to rely more on lipolysis and fatty acid metabolism, while males preferentially use carbohydrates (Hedrington and Davis, 2015; Klosinski et al., 2015; Yao et al., 2010). The distribution of energy stores and the patterns of energy utilization during weight loss are also sexually dimorphic (Palmer and Clegg, 2015; Pietrobelli et al., 2002), and when considered with the recent finding that AgRP neurons regulate substrate utilization in metabolism and lipogenesis (Cavalcanti-de-Albuquerque et al., 2019), this sex-dependent difference in AgRP neuronal function may reflect one mechanism by which females regulate metabolic flexibility.

We did not investigate the impact of HFD on female body weight, body composition, or neuronal function past 8 weeks of HFD feeding and all of the mice used here were less than 6 months old and undergoing normal estrous cycles. Estrogen is an important determinant of feeding and diet-induced increases in adiposity (Litwak et al., 2014), possibly in part via actions on AgRP neurons (Olofsson et al., 2009). Thus, female mice on HFD for >8 weeks may undergo age- and diet-induced changes in AgRP neuronal function and body weight more similar to what we have observed in young male mice. The impact of longer (>8 weeks) HFD feeding on AgRP neuronal function and leptin sensitivity in female mice will be determined in future studies.

### HFD decouples the intrinsic excitability of AgRP neurons from physiological and synaptic cues that match neuronal function to metabolic state

We previously demonstrate that HFD feeding was associated with plasticity in the intrinsic excitability of AgRP neurons and that these neurons become resistant to inhibition by leptin (Baver et al., 2014). In addition of hormonal modulation, AgRP neurons have been shown to be modulated by synaptic inputs – increased excitatory glutamatergic input involving the action of NMDA receptors is required for the fasting-induced increase in AgRP neuronal activity (Liu et al., 2012; Yang et al., 2011) and more recently, several groups have used *in vivo* approaches to demonstrate that the sensory detection of food rapidly and robustly inhibits AgRP neurons in a manner that does not require the ingestion of food (Betley et al., 2015; Chen et al., 2015; Mandelblat-Cerf et al., 2015). Thus, the activity of AgRP neurons is highly regulated by synaptic input from upstream (largely unknown) circuits, which may represent an additional target for diet-induced plasticity. We investigated this possibility using whole-cell patch clamp recording of synaptic activity to dissect the effects of HFD and obesity on both excitatory and inhibitory synaptic input to AgRP neurons.

Excitatory glutamatergic input is required for the fasting-induced increase in AgRP neuronal excitability (Liu et al., 2012; Yang et al., 2011), we wanted to determine if a similar phenomenon was associated with the increase in AgRP neuronal excitability after brief periods of HFD feeding (Wei et al., 2015). Short (2d) periods of HFD feeding were associated with an increase in the frequency of excitatory input to AgRP neurons, with no change in the rate of inhibitory input, suggesting an imbalance in the excitation/inhibition (E/I) ratio arising from the presentation and/or consumption of a novel high-fat food (Figure 3). This finding in consistent with a recent study using *in vivo* fiber photometry to monitor activity in AgRP neurons from mice fed HFD (Mazzone et al., 2020). In this study, the inhibitory response of AgRP neurons to the presentation of food was significantly blunted in mice fed a HFD for only 1 week but exhibited appropriate responses to gut-derived satiety signals such as cholecystokinin or peptide PYY, suggesting that this early response to HFD may arise from a transient increase in excitatory input rather than obesity-induced metabolic dysfunction. Interestingly, after 2d of HFD, AgRP neurons from female mice exhibited a significant decrease in inhibitory GABAergic input (Figure 3E) rather than an increase in glutamatergic input (Figure 3B), suggesting that sex may play an important role in the modulation of AgRP neurons in response to food.

This increase in excitatory synaptic input to AgRP neurons is transient and by 8 weeks of HFD feeding, the baseline level of glutamatergic input to AgRP neurons has returned to baseline values (Figure 4C). However, we do not believe this decrease represents a diet-induced change in presynaptic input to AgRP neurons, as neurons from both male and female HFD mice respond to fasting similarly to those from mice maintained on NCD (Figure 4, C and D), suggesting that the presynaptic neurons involved in this response remain sensitive and respond appropriately to changes in the animal’s metabolic state. However, despite this seemingly normal presynaptic response to food deprivation, the intrinsic excitability of the postsynaptic AgRP neuron is no longer coupled to alterations in synaptic input after HFD feeding, as the spontaneous firing rate of AgRP neurons from HFD_fed_ and HFD_fast_ mice is nearly identical (Baver et al., 2014), indicating that following long-term HFD feeding, AgRP neurons become desensitized to excitatory synaptic input. We see a similar response in AgRP neurons from female mice, who do not exhibit diet-induced obesity or leptin resistance, yet still appear to have become desensitized to excitatory input. It will be important to determine if this reflects a common mechanism that is independent of obesity and leptin signaling or if this response is sex-dependent.

### Long-term HFD consumption disrupts GABAergic inhibition by inducing dyshomeostasis of intracellular Cl-in AgRP neurons

Neuronal excitability is tightly governed by the coordination of excitatory and inhibitory inputs (He and Cline, 2019). Given that we found HFD had a minimal impact on excitatory inputs to AgRP neurons, we reasoned that perhaps diet-induced hyperexcitability of AgRP neurons was due to a loss of inhibitory synaptic input, but surprisingly we found the opposite was true – HFD diet is associated with a significant increase in GABAergic input, a finding that was unexpected given the diet-induced hyperexcitability of AgRP neurons. We confirmed that the defect in GABAergic signaling is likely to be entirely due to postsynaptic effects, as AgRP neurons from HFD mice were not inhibited by either the direct application of GABA or by the optogenetic activation of the GABAergic vDMH→ARH^AgRP^ circuit (Figure 6); in fact, a subset of neurons from HFD mice were excited by GABA (Figure 6D), indicating that intracellular Cl^−^, which is the major determinant of GABA-mediated inhibition via GABA_A_-R, is significantly disrupted following long-term HFD feeding. We confirmed this by directly measuring E_GABA_, which is directly related to intracellular Cl^−^ concentration, and found that E_GABA_ is significantly depolarized in AgRP neurons from HFD_8w_ mice, providing a mechanistic explanation for the loss of GABAergic inhibition.

Based on these results, we propose that dysfunction of the neuronal K^+^/Cl^−^ cotransporter KCC2 may underlie many of the observed effects of HFD on AgRP neuronal excitability. KCC2 is the primary Cl^−^ extruder in mature neurons, including AgRP neurons and under normal conditions, is responsible for keeping intraneuronal Cl^−^ low, thus ensuring that E_GABA_ is hyperpolarized relative to the resting membrane potential and forming the biophysical basis for the inhibitory action of GABA_A_-R. The importance of this function is perhaps best exemplified by the switch of GABA from excitatory to inhibitory subsequent to upregulation of KCC2 during development, however, it is increasingly apparent that KCC2 dysfunction is a contributing factor in several adult-onset disorders, including epilepsy (Moore et al., 2017, 2018), ischemic stroke (Jaenisch Nadine et al., 2010), Huntington’s disease (Dargaei et al., 2018), and peripheral nerve injury (Boulenguez et al., 2010; Chen et al., 2018). Our finding that GABA and the activation of GABAergic terminals no longer inhibit AgRP neurons and that E_GABA_ is significantly depolarized in AgRP neurons following prolonged HFD feeding strongly implicate KCC2 dysfunction as a causal factor in obesity.

KCC2 is an attractive candidate for the diet-induced synaptic dysfunction and hyperexcitability of AgRP neurons from HFD fed mice for several additional reasons. Beyond its canonical role as the primary Cl^−^ extruder, KCC2 has been shown to additionally play a critical role in the maintenance and function of excitatory synapses, including the maturation of dendritic spines, trafficking and retention of AMPA receptors, and induction of LTP (Chamma et al., 2012, 2013; Chevy et al., 2015; Gauvain et al., 2011). Elegant proteomics experiments have revealed that KCC2 has a large, diverse interactome (Mahadevan et al., 2017; Smalley et al., 2020) and that more than 40% of its interaction partners are associated with excitatory synapses (Mahadevan et al., 2017; Pressey et al., 2020). Diet-induced KCC2 dysfunction may therefore also play a causal or contributing role to the dysfunction we found in postsynaptic glutamatergic signaling in AgRP neurons.

Lastly, KCC2 likely functions as a hub protein within its large interactome (Pressey et al., 2020), therefore dysfunction in the expression, function or membrane targeting of KCC2 is predicted to have wide-ranging impact on neuronal excitability. For this reason and those outlined above, diet-induced dysfunction of KCC2 is a parsimonious explanation for the remodeling of AgRP neurons by HFD and represents a mechanistic explanation for the changes in AgRP neuronal function reported by us and others (Beutler et al., 2020; Mazzone et al., 2020). Therapeutic interventions that aim to restore the normal function of this critical protein may represent a much-needed tool for the treatment of obesity and its comorbidities.

## References

Abdelaal, M., le Roux, C.W., and Docherty, N.G. (2017). Morbidity and mortality associated with obesity. Ann. Transl. Med. 5.

Andrews, Z.B., Liu, Z.-W., Walllingford, N., Erion, D.M., Borok, E., Friedman, J.M., Tschöp, M.H., Shanabrough, M., Cline, G., Shulman, G.I., et al. (2008). UCP2 mediates ghrelin’s action on NPY/AgRP neurons by lowering free radicals. Nature 454, 846–851.

Aponte, Y., Atasoy, D., and Sternson, S.M. (2011). AGRP neurons are sufficient to orchestrate feeding behavior rapidly and without training. Nat. Neurosci. 14, 351–355.

Atamni, H.J.A.-T., Mott, R., Soller, M., and Iraqi, F.A. (2016). High-fat-diet induced development of increased fasting glucose levels and impaired response to intraperitoneal glucose challenge in the collaborative cross mouse genetic reference population. BMC Genet. 17, 10.

Baver, S.B., Hope, K., Guyot, S., Bjørbaek, C., Kaczorowski, C., and O’Connell, K.M.S. (2014). Leptin Modulates the Intrinsic Excitability of AgRP/NPY Neurons in the Arcuate Nucleus of the Hypothalamus. J. Neurosci. 34, 5486–5496.

Berk, K.A., Buijks, H.I.M., Verhoeven, A.J.M., Mulder, M.T., Özcan, B., van’t Spijker, A., Timman, R., Busschbach, J.J., and Sijbrands, E.J. (2018). Group cognitive behavioural therapy and weight regain after diet in type 2 diabetes: results from the randomised controlled POWER trial. Diabetologia 61, 790–799.

Betley, J.N., Xu, S., Cao, Z.F.H., Gong, R., Magnus, C.J., Yu, Y., and Sternson, S.M. (2015). Neurons for hunger and thirst transmit a negative-valence teaching signal. Nature 521, 180–185.

Beutler, L.R., Corpuz, T.V., Ahn, J.S., Kosar, S., Song, W., Chen, Y., and Knight, Z.A. (2020). Obesity causes selective and long-lasting desensitization of AgRP neurons to dietary fat. ELife 9, e55909.

Boulenguez, P., Liabeuf, S., Bos, R., Bras, H., Jean-Xavier, C., Brocard, C., Stil, A., Darbon, P., Cattaert, D., Delpire, E., et al. (2010). Down-regulation of the potassium-chloride cotransporter KCC2 contributes to spasticity after spinal cord injury. Nat. Med. 16, 302–307.

Cavalcanti-de-Albuquerque, J.P., Bober, J., Zimmer, M.R., and Dietrich, M.O. (2019). Regulation of substrate utilization and adiposity by Agrp neurons. Nat. Commun. 10, 311.

Chamma, I., Chevy, Q., Poncer, J.C., and Levi, S. (2012). Role of the neuronal K-Cl co-transporter KCC2 in inhibitory and excitatory neurotransmission. Front. Cell. Neurosci. 6.

Chamma, I., Heubl, M., Chevy, Q., Renner, M., Moutkine, I., Eugène, E., Poncer, J.C., and Lévi, S. (2013). Activity-dependent regulation of the K/Cl transporter KCC2 membrane diffusion, clustering, and function in hippocampal neurons. J. Neurosci. Off. J. Soc. Neurosci. 33, 15488–15503.

Chen, B., Li, Y., Yu, B., Zhang, Z., Brommer, B., Williams, P.R., Liu, Y., Hegarty, S.V., Zhou, S., Zhu, J., et al. (2018). Reactivation of Dormant Relay Pathways in Injured Spinal Cord by KCC2 Manipulations. Cell 174, 521–535.e13.

Chen, Y., Lin, Y.-C., Kuo, T.-W., and Knight, Z.A. (2015). Sensory Detection of Food Rapidly Modulates Arcuate Feeding Circuits. Cell 160, 829–841.

Chen, Y., Lin, Y.-C., Zimmerman, C.A., Essner, R.A., and Knight, Z.A. (2016). Hunger neurons drive feeding through a sustained, positive reinforcement signal. ELife 5, e18640.

Chevy, Q., Heubl, M., Goutierre, M., Backer, S., Moutkine, I., Eugène, E., Bloch-Gallego, E., Lévi, S., and Poncer, J.C. (2015). KCC2 Gates Activity-Driven AMPA Receptor Traffic through Cofilin Phosphorylation. J. Neurosci. Off. J. Soc. Neurosci. 35, 15772–15786.

Dargaei, Z., Bang, J.Y., Mahadevan, V., Khademullah, C.S., Bedard, S., Parfitt, G.M., Kim, J.C., and Woodin, M.A. (2018). Restoring GABAergic inhibition rescues memory deficits in a Huntington’s disease mouse model. Proc. Natl. Acad. Sci. U. S. A. 115, E1618–E1626.

Dorfman, M.D., Krull, J.E., Douglass, J.D., Fasnacht, R., Lara-Lince, F., Meek, T.H., Shi, X., Damian, V., Nguyen, H.T., Matsen, M.E., et al. (2017). Sex differences in microglial CX3CR1 signalling determine obesity susceptibility in mice. Nat. Commun. 8, 14556.

Egan, O.K., Inglis, M.A., and Anderson, G.M. (2017). Leptin Signaling in AgRP Neurons Modulates Puberty Onset and Adult Fertility in Mice. J. Neurosci. 37, 3875–3886.

El-Haschimi, K., Pierroz, D.D., Hileman, S.M., Bjørbæk, C., and Flier, J.S. (2000). Two defects contribute to hypothalamic leptin resistance in mice with diet-induced obesity. J. Clin. Invest. 105, 1827–1832.

Foster, G.D., Wyatt, H.R., Hill, J.O., Makris, A.P., Rosenbaum, D.L., Brill, C., Stein, R.I., Mohammed, B.S., Miller, B., Rader, D.J., et al. (2010). Weight and Metabolic Outcomes After 2 Years on a Low-Carbohydrate Versus Low-Fat Diet. Ann. Intern. Med. 153, 147–157.

Fothergill, E., Guo, J., Howard, L., Kerns, J.C., Knuth, N.D., Brychta, R., Chen, K.Y., Skarulis, M.C., Walter, M., Walter, P.J., et al. (2016). Persistent metabolic adaptation 6 years after “The Biggest Loser” competition. Obes. Silver Spring Md 24, 1612–1619.

Gao, F., Zheng, K.I., Wang, X.-B., Sun, Q.-F., Pan, K.-H., Wang, T.-Y., Chen, Y.-P., Targher, G., Byrne, C.D., George, J., et al. (2020). Obesity Is a Risk Factor for Greater COVID-19 Severity. Diabetes Care 43, e72–e74.

Garfield, A.S., Shah, B.P., Burgess, C.R., Li, M.M., Li, C., Steger, J.S., Madara, J.C., Campbell, J.N., Kroeger, D., Scammell, T.E., et al. (2016). Dynamic GABAergic afferent modulation of AgRP neurons. Nat. Neurosci. 19, 1628–1635.

Gauvain, G., Chamma, I., Chevy, Q., Cabezas, C., Irinopoulou, T., Bodrug, N., Carnaud, M., Lévi, S., and Poncer, J.C. (2011). The neuronal K-Cl cotransporter KCC2 influences postsynaptic AMPA receptor content and lateral diffusion in dendritic spines. Proc. Natl. Acad. Sci. U. S. A. 108, 15474–15479.

Greenway, F.L. (2015). Physiological adaptations to weight loss and factors favouring weight regain. Int. J. Obes. 39, 1188–1196.

Hales, C.M. (2020). Prevalence of Obesity and Severe Obesity Among Adults: United States, 2017–2018. 8.

Hall, K.D., and Guo, J. (2017). Obesity Energetics: Body Weight Regulation and the Effects of Diet Composition. Gastroenterology 152, 1718–1727.e3.

He, H., and Cline, H.T. (2019). What Is Excitation/Inhibition and How Is It Regulated? A Case of the Elephant and the Wisemen. J. Exp. Neurosci. 13.

Hedrington, M.S., and Davis, S.N. (2015). Sexual Dimorphism in Glucose and Lipid Metabolism during Fasting, Hypoglycemia, and Exercise. Front. Endocrinol. 6.

Hong, J., Stubbins, R.E., Smith, R.R., Harvey, A.E., and Núñez, N.P. (2009). Differential susceptibility to obesity between male, female and ovariectomized female mice. Nutr. J. 8, 11.

Hwang, L.-L., Wang, C.-H., Li, T.-L., Chang, S.-D., Lin, L.-C., Chen, C.-P., Chen, C.-T., Liang, K.-C., Ho, I.-K., Yang, W.-S., et al. (2010). Sex Differences in High-fat Diet-induced Obesity, Metabolic Alterations and Learning, and Synaptic Plasticity Deficits in Mice. Obesity 18, 463–469.

Jaenisch Nadine, Witte Otto W., and Frahm Christiane (2010). Downregulation of Potassium Chloride Cotransporter KCC2 After Transient Focal Cerebral Ischemia. Stroke 41, e151–e159.

Klosinski, L.P., Yao, J., Yin, F., Fonteh, A.N., Harrington, M.G., Christensen, T.A., Trushina, E., and Brinton, R.D. (2015). White Matter Lipids as a Ketogenic Fuel Supply in Aging Female Brain: Implications for Alzheimer’s Disease. EBioMedicine 2, 1888–1904.

Könner, A.C., Janoschek, R., Plum, L., Jordan, S.D., Rother, E., Ma, X., Xu, C., Enriori, P., Hampel, B., Barsh, G.S., et al. (2007). Insulin Action in AgRP-Expressing Neurons Is Required for Suppression of Hepatic Glucose Production. Cell Metab. 5, 438–449.

Krashes, M.J., Koda, S., Ye, C., Rogan, S.C., Adams, A.C., Cusher, D.S., Maratos-Flier, E., Roth, B.L., and Lowell, B.B. (2011). Rapid, reversible activation of AgRP neurons drives feeding behavior in mice. J. Clin. Invest. 121, 1424–1428.

Krashes, M.J., Shah, B.P., Madara, J.C., Olson, D.P., Strochlic, D.E., Garfield, A.S., Vong, L., Pei, H., Watabe-Uchida, M., Uchida, N., et al. (2014). An excitatory paraventricular nucleus to AgRP neuron circuit that drives hunger. Nature 507, 238–242.

Kyle, T.K., Dhurandhar, E.J., and Allison, D.B. (2016). Regarding Obesity as a Disease: Evolving Policies and Their Implications. Endocrinol. Metab. Clin. North Am. 45, 511–520.

Li, X.-Y., Han, Y., Zhang, W., Wang, S.-R., Wei, Y.-C., Li, S.-S., Lin, J.-K., Yan, J.-J., Chen, A.-X., Zhang, X., et al. (2019). AGRP Neurons Project to the Medial Preoptic Area and Modulate Maternal Nest-Building. J. Neurosci. 39, 456–471.

Litwak, S.A., Wilson, J.L., Chen, W., Garcia-Rudaz, C., Khaksari, M., Cowley, M.A., and Enriori, P.J. (2014). Estradiol Prevents Fat Accumulation and Overcomes Leptin Resistance in Female High-Fat Diet Mice. Endocrinology 155, 4447–4460.

Liu, T., Kong, D., Shah, B.P., Ye, C., Koda, S., Saunders, A., Ding, J.B., Yang, Z., Sabatini, B.L., and Lowell, B.B. (2012). Fasting Activation of AgRP Neurons Requires NMDA Receptors and Involves Spinogenesis and Increased Excitatory Tone. Neuron 73, 511–522.

Luquet, S., Perez, F.A., Hnasko, T.S., and Palmiter, R.D. (2005). NPY/AgRP Neurons Are Essential for Feeding in Adult Mice but Can Be Ablated in Neonates. Science 310, 683–685.

Mahadevan, V., Khademullah, C.S., Dargaei, Z., Chevrier, J., Uvarov, P., Kwan, J., Bagshaw, R.D., Pawson, T., Emili, A., De Koninck, Y., et al. (2017). Native KCC2 interactome reveals PACSIN1 as a critical regulator of synaptic inhibition. ELife 6.

Mandelblat-Cerf, Y., Ramesh, R.N., Burgess, C.R., Patella, P., Yang, Z., Lowell, B.B., and Andermann, M.L. (2015). Arcuate hypothalamic AgRP and putative POMC neurons show opposite changes in spiking across multiple timescales. ELife 4, e07122.

Masters, R.K., Reither, E.N., Powers, D.A., Yang, Y.C., Burger, A.E., and Link, B.G. (2013). The Impact of Obesity on US Mortality Levels: The Importance of Age and Cohort Factors in Population Estimates. Am. J. Public Health 103, 1895–1901.

Mazzone, C.M., Liang-Guallpa, J., Li, C., Wolcott, N.S., Boone, M.H., Southern, M., Kobzar, N.P., Salgado, I. de A., Reddy, D.M., Sun, F., et al. (2020). High-fat food biases hypothalamic and mesolimbic expression of consummatory drives. Nat. Neurosci. 23, 1253–1266.

Moore, Y.E., Kelley, M.R., Brandon, N.J., Deeb, T.Z., and Moss, S.J. (2017). Seizing Control of KCC2: A New Therapeutic Target for Epilepsy. Trends Neurosci. 40, 555–571.

Moore, Y.E., Deeb, T.Z., Chadchankar, H., Brandon, N.J., and Moss, S.J. (2018). Potentiating KCC2 activity is sufficient to limit the onset and severity of seizures. Proc. Natl. Acad. Sci. U. S. A. 115, 10166–10171.

Morton, G.J., and Schwartz, M.W. (2001). The NPY/AgRP neuron and energy homeostasis. Int. J. Obes. 25, S56–S62.

Münzberg, H., Flier, J.S., and Bjørbæk, C. (2004). Region-Specific Leptin Resistance within the Hypothalamus of Diet-Induced Obese Mice. Endocrinology 145, 4880–4889.

Nordmo, M., Danielsen, Y.S., and Nordmo, M. (2020). The challenge of keeping it off, a descriptive systematic review of high-quality, follow-up studies of obesity treatments. Obes. Rev. 21, e12949.

Olofsson, L.E., Pierce, A.A., and Xu, A.W. (2009). Functional requirement of AgRP and NPY neurons in ovarian cycle-dependent regulation of food intake. Proc. Natl. Acad. Sci. 106, 15932–15937.

Padilla, S.L., Qiu, J., Nestor, C.C., Zhang, C., Smith, A.W., Whiddon, B.B., Rønnekleiv, O.K., Kelly, M.J., and Palmiter, R.D. (2017). AgRP to Kiss1 neuron signaling links nutritional state and fertility. Proc. Natl. Acad. Sci. 114, 2413–2418.

Palmer, B.F., and Clegg, D.J. (2015). The sexual dimorphism of obesity. Mol. Cell. Endocrinol. 402, 113–119.

Perez-Sieira, S., Martinez, G., Porteiro, B., Lopez, M., Vidal, A., Nogueiras, R., and Dieguez, C. (2013). Female Nur77-deficient mice show increased susceptibility to diet-induced obesity. PloS One 8, e53836.

Pettersson, U.S., Waldén, T.B., Carlsson, P.-O., Jansson, L., and Phillipson, M. (2012). Female Mice are Protected against High-Fat Diet Induced Metabolic Syndrome and Increase the Regulatory T Cell Population in Adipose Tissue. PLOS ONE 7, e46057.

Phillips, C.T., and Palmiter, R.D. (2008). Role of Agouti-Related Protein-Expressing Neurons in Lactation. Endocrinology 149, 544–550.

Pietrobelli, A., Allison, D.B., Heshka, S., Heo, M., Wang, Z.M., Bertkau, A., Laferrère, B., Rosenbaum, M., Aloia, J.F., Pi-Sunyer, F.X., et al. (2002). Sexual dimorphism in the energy content of weight change. Int. J. Obes. 26, 1339–1348.

Popkin, B.M., Du, S., Green, W.D., Beck, M.A., Algaith, T., Herbst, C.H., Alsukait, R.F., Alluhidan, M., Alazemi, N., and Shekar, M. (2020). Individuals with obesity and COVID-19: A global perspective on the epidemiology and biological relationships. Obes. Rev. 21, e13128.

Pressey, J.C., Mahadevan, V., and Woodin, M.A. (2020). Chapter 8 - KCC2 is a hub protein that balances excitation and inhibition. In Neuronal Chloride Transporters in Health and Disease, X. Tang, ed. (Academic Press), pp. 159–179.

Profenno, L.A., Porsteinsson, A.P., and Faraone, S.V. (2010). Meta-Analysis of Alzheimer’s Disease Risk with Obesity, Diabetes, and Related Disorders. Biol. Psychiatry 67, 505–512.

Pugazhenthi, S., Qin, L., and Reddy, P.H. (2017). Common neurodegenerative pathways in obesity, diabetes, and Alzheimer’s disease. Biochim. Biophys. Acta Mol. Basis Dis. 1863, 1037–1045.

Rossi, M.A., Basiri, M.L., McHenry, J.A., Kosyk, O., Otis, J.M., Munkhof, H.E. van den, Bryois, J., Hübel, C., Breen, G., Guo, W., et al. (2019). Obesity remodels activity and transcriptional state of a lateral hypothalamic brake on feeding. Science 364, 1271–1274.

Salinero, A.E., Anderson, B.M., and Zuloaga, K.L. (2018). Sex differences in the metabolic effects of diet-induced obesity vary by age of onset. Int. J. Obes. 42, 1088–1091.

Smalley, J.L., Kontou, G., Choi, C., Ren, Q., Albrecht, D., Abiraman, K., Santos, M.A.R., Bope, C.E., Deeb, T.Z., Davies, P.A., et al. (2020). Isolation and Characterization of Multi-Protein Complexes Enriched in the K-Cl Co-transporter 2 From Brain Plasma Membranes. Front. Mol. Neurosci. 13.

Spring, S., Lerch, J.P., and Henkelman, R.M. (2007). Sexual dimorphism revealed in the structure of the mouse brain using three-dimensional magnetic resonance imaging. NeuroImage 35, 1424–1433.

Sumithran, P., and Proietto, J. (2013). The defence of body weight: A physiological basis: For weight regain after weight loss. Clin. Sci. Lond. Engl. 1979 124, 231–241.

Takahashi, K.A., and Cone, R.D. (2005). Fasting induces a large, leptin-dependent increase in the intrinsic action potential frequency of orexigenic arcuate nucleus neuropeptide Y/Agouti-related protein neurons. Endocrinology 146, 1043–1047.

van den Top, M., Lee, K., Whyment, A.D., Blanks, A.M., and Spanswick, D. (2004). Orexigen-sensitive NPY/AgRP pacemaker neurons in the hypothalamic arcuate nucleus. Nat. Neurosci. 7, 493–494.

Van der Ploeg, L.H. (2000). Obesity: an epidemic in need of therapeutics. Curr. Opin. Chem. Biol. 4, 452–460.

Varela, L., and Horvath, T.L. (2012). Leptin and insulin pathways in POMC and AgRP neurons that modulate energy balance and glucose homeostasis. EMBO Rep. 13, 1079–1086.

Wang, C., Zhou, W., He, Y., Yang, T., Xu, P., Yang, Y., Cai, X., Wang, J., Liu, H., Yu, M., et al. (2021). AgRP neurons trigger long-term potentiation and facilitate food seeking. Transl. Psychiatry 11, 1–17.

Webb, V.L., and Wadden, T.A. (2017). Intensive Lifestyle Intervention for Obesity: Principles, Practices, and Results. Gastroenterology 152, 1752–1764.

Wei, W., Pham, K., Gammons, J.W., Sutherland, D., Liu, Y., Smith, A., Kaczorowski, C.C., and O’Connell, K.M.S. (2015). Diet composition, not calorie intake, rapidly alters intrinsic excitability of hypothalamic AgRP/NPY neurons in mice. Sci. Rep. 5, 16810.

Yakar, S., Nunez, N.P., Pennisi, P., Brodt, P., Sun, H., Fallavollita, L., Zhao, H., Scavo, L., Novosyadlyy, R., Kurshan, N., et al. (2006). Increased tumor growth in mice with diet-induced obesity: impact of ovarian hormones. Endocrinology 147, 5826–5834.

Yang, Y., Atasoy, D., Su, H.H., and Sternson, S.M. (2011). Hunger States Switch a Flip-Flop Memory Circuit via a Synaptic AMPK-Dependent Positive Feedback Loop. Cell 146, 992–1003.

Yao, J., Hamilton, R.T., Cadenas, E., and Brinton, R.D. (2010). Decline in mitochondrial bioenergetics and shift to ketogenic profile in brain during reproductive senescence. Biochim. Biophys. Acta 1800, 1121–1126.

